# Phenotypic spandrel: absolute discrimination and ligand antagonism

**DOI:** 10.1101/036293

**Authors:** Paul Francois, Kyle A Johnson, Laura N Saunders

**Affiliations:** Physics Department, McGill University, Montreal, Quebec, Canada H3A 2T8

## Abstract

We consider the general problem of absolute discrimination between categories of ligands irrespective of their concentration. An instance of this problem is immune discrimination between self and not-self. We connect this problem to biochemical adaptation, and establish that ligand antagonism - the ability of sub threshold ligands to negatively impact response - is a necessary consequence of absolute discrimination.Thus antagonism constitutes a “phenotypic spandrel”: a phenotype existing as a necessary by-product of another phenotype. We exhibit a simple analytic model of absolute discrimination displaying ligand antagonism, where antagonism strength is linear in distance from threshold. This contrasts with proofreading based models, where antagonism vanishes far from threshold and thus displays an inverted hierarchy of antagonism compared to simple model. The phenotypic spandrel studied here is expected to structure many decision pathways such as immune detection mediated by TCRs and FceRIs.

---

Recent works in quantitative evolution combined to mathematical modelling have shown that evolution of biological structures is constrained by selected phenotypes in strong unexpected ways. Trade-off between different functionalities are major forces shaping evolution of complex phenotypes moving on evolutionary Pareto fronts [1, 2]. *In silico* evolution of phenotypic models of gene networks [3] have further shown that selection of complex phenotypes leads to apparition of other traits that have not been explicitly selected for. For instance, a clock naturally appears when selecting for stripe formation in a model of segmentation evolution [4], or ligand antagonism when selecting for a model of immune recognition [5]. This is reminiscent of the architectural image of “evolutionary spandrels” proposed by Gould and Lewontin [6]. They argued that many biological properties are necessary by-products of more fundamental adaptive traits, that can be further elaborated by evolution (leading to the notion of “exaptation”[7]).

An important biological example is the absolute discrimination between different ligand “qualities”, an instance being early immune recognition [8–11]. T cells have to recognize minute concentrations of not-self while being insensitive to overwhelming concentration of self ligands. The parameter discriminating self from not-self is the ligand dissociation time *τ* [12, 13] to T cell receptors (TCRs), defining the so-called “life-time dogma” [9]. In evolutionary simulations [5], the phenomenon of absolute discrimination is not achieved without detrimental ligand antagonism: an experimental “dog in the manger” effect in which ligands unable to trigger response prevent agonists to do so [14, 15].

Here, we show mathematically that ligand antagonism is a necessary by-product of absolute discrimination. We thus qualify antagonism as a “phenotypic spandrel”: a phenotype existing as a necessary by-product of another phenotype. We exhibit a generic simple model for antagonism, and further show how addition of proofreading steps leads to an “inverted” hierarchy of antagonism that is *not* a generic feature of the antagonism spandrel.

We consider a family of biochemical ligands, with different quantitative properties encoded by a continuous parameter *τ*, subsequently called “ligand quality”. Ideal absolute discrimination is observed when cells successfully discriminate between two categories of ligands, above and below critical quality *τ_c_*, irrespective of ligand quantity *L*. An idealized response diagram for absolute discrimination in (*L*, *τ*) plane defines a vertical “discrimination line” at *τ_c_* (Fig1 A).

**FIG. 1.**
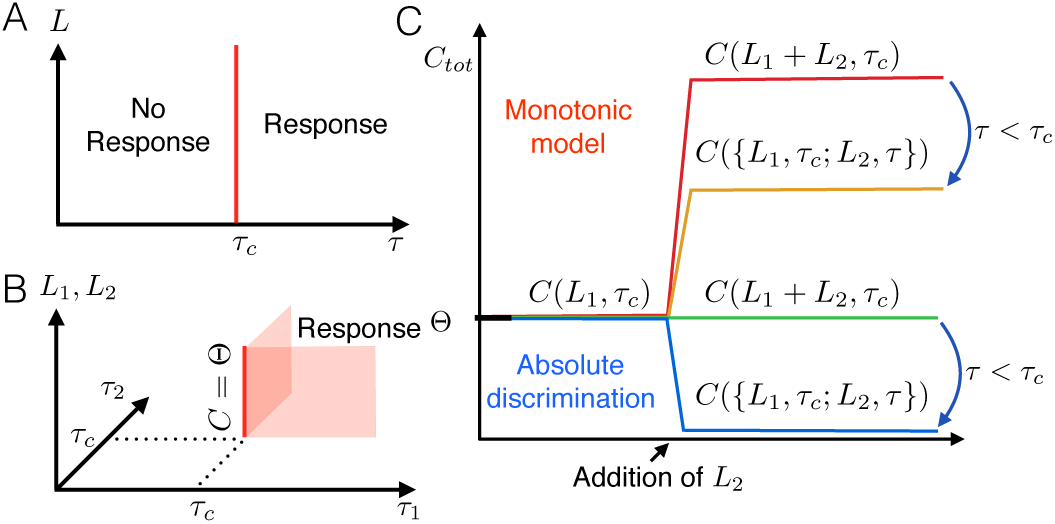
(A). Sketch of absolute discrimination for a single ligand type in (*L, τ*) plane. Discrimination line corresponding to *τ = τ_c_* is in red. (B). Sketch of absolute discrimination for mixtures, adding a second type of ligands with quality *τ_2_*, with *L_1_, L_2_* ligand concentrations represented by a vertical axis. The discrimination line for *τ*_1_ = *τ*_2_ = *τ_c_* is vertical and satisfies equation 1. Red planes correspond to mixtures of agonist where necessary 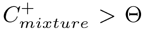. (C). Intuitive explanation of antagonism (see main text). Relative height of mixtures correspond to values of *C*. For absolute discrimination, any amount of ligand at *τ_c_* is on the discrimination line, so that if *τ < τ_c_*, contribution of *L*_2_ ligands is negative.

We model absolute discrimination using biochemical networks with continuous, single-valued variables, at steady state, in a way similar to most phenotypic models of early immune recognition such as the ones reviewed in [10]. Discrimination models are often based on comparison of a downstream protein *C* concentration to a threshold Θ, i.e. response is triggered when *C ≥* Θ. For ligands at *τ = τ_c_* from Figure 1A, by continuity we thus necessarily have on the discrimination line

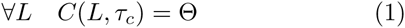

Equation 1 expresses that *C* is independent from the concentration of ligand *L:* technically *C* is a biochemically adaptive variable at *τ = τ_c_* [16, 17].

*In vivo*, cells are exposed to complex mixtures of ligands interacting with receptors at the surface of a single cell. Consider now the problem of absolute discrimination in mixtures, sketched on Fig 1B. Assume that on the discrimination line *τ = τ_c_*, we change the quality of a fraction *r* of ligands from 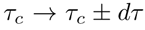. Assuming for *dτ* > 0 a mixture of (1−*r)L* ligands at *τ_c_* with *rL* ligands at *τ_c_ + dτ* yields response so that for the corresponding output variable

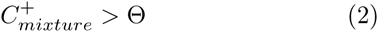

We can then Taylor expand this expression with respect to *dτ* to define

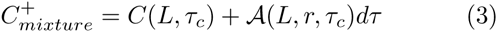

and then combining with equations 1-2 we get *A*(*L, r, τ_c_*) > 0. Flipping the sign in front of *dτ* to consider another type of ligand mixture (1 - *r)L* ligands at *T_C_* and *rL* ligands at *τ_c_* − *dτ*, we thus have

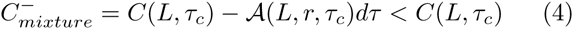

This inequality is valid for any *r*, *L*. We then change our variables to define the mixture as *L_1_ = L(1 - r*) agonist ligands mixed with *L_2_ = rL* sub threshold ligands (i.e. *r = L_2_*/(*L_1_ + L_2_*),*L = L_1_ + L_2_*). Furthermore from equation 1 for any r value we have *C*(*L, τ_c_*) = *C*((1 −*r*)*L, τ_c_*) = *Θ*, so substituting in right-hand side of 4 we get:

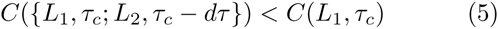

This latter expression is the exact definition of antagonism: as soon as sub-threshold ligands *L_2_* are added in presence of critical agonist ligands *L*_1_, the output variable is decreased with respect to the case of pure *L*_1_, and thus if decision is based on thresholding of output, response disappears.

An intuitive explanation of the above reasoning can be made. Considering a mixture of ligands (*L_n_*) of different qualities (*τ_n_*), output *C* is an increasing function of both *L* and *τ* for models with monotonic response curves such as kinetic proofreading [18]. In such models, starting from a critical ligand concentration *L_1_* triggering response (i.e. *C*(*L*_1_, *τ_c_*) = *Θ*), any addition of ligands *L_2_* gives higher output value and response is maintained. Addition of *L_2_* ligands with critical quality *τ_c_* gives higher output than addition of same quantity *L_2_* of ligands with lower quality *τ < τ_c_* (symbolized by a downward arrow in the upper part of Figure 1C).

Considering now absolute discrimination and again addition of *L_2_* ligands on Figure 1C, the constraint encoded by equation 1 means that addition of extra *L_2_* ligands of quality *τ = τ_c_* does not impact output value. But if we require that mixtures of agonists trigger response (expressed by equation 2), it means that output sill is a monotonic function of *τ*. Thus if ligand quality of the extra *L_2_* ligands is lowered, we expect that a negative contribution similar to the monotonic example (with *τ < τ_c_*) appears (symbolized by a downward arrow in the lower part of Figure 1C). Thus response would now be below threshold of activation, corresponding to antagonism.

Antagonism here is a direct consequence of biochemical adaptation at *τ* = *τ_c_*. By continuity, antagonism is expected for ligands not necessarily tuned on the critical line *τ = τ_c_*. Furthermore, adaptation does not necessarily have to be perfect: non-monotonic response curves [19], with *C* varying around Θ, essentially approximate well adaptation necessary for absolute discrimination and display antagonism [5, 20]. At least two examples of absolute discrimination/antagonism are offered in the immune context. For early immune recognition mediated by TCRs, agonists simultaneously presented with ligands just below threshold fail to trigger response, while agonists alone do [15, 21]. For FceRIs mediated response in mast cells [14], a very similar “dog in the manger” effect is observed where low affinity antagonists titrate the Lyn kinase responsible for proofreading steps (exactly like adaptive sorting evolved in [5]).

Further categorization of absolute discrimination mechanisms combined to antagonism is possible. A “homeostatic model” (Figure 2) is inspired by a ligand-receptor adaptive network evolved in [16].

**FIG. 2.**
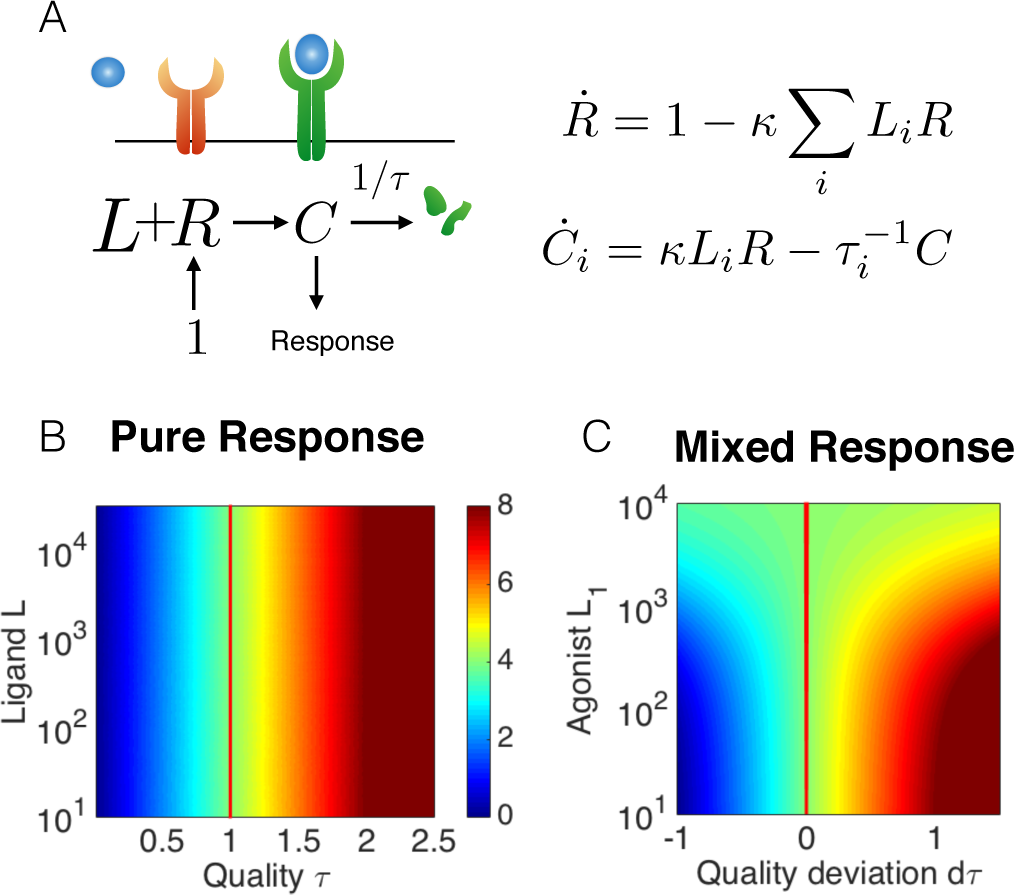
(A). Sketch of homeostatic network considered in main text, with corresponding equations. For simulations we use κ = 10^−3^, and define *τ_c_* = Θ = 4. (B). Color map of output *C* values in the (*L, τ*) plane, ligand quality is plotted in units of *τ_c_*.(B) Discrimination line (solid red) is vertical similar to figure 1 A (C) Color map of output *C* values in plane (L1, *dτ*) for mixtures of L2 = 10 ligands, quality *τ*+*dτ*, with *L1* ligands, quality *τ_c_*. This plane is orthogonal to the plane displayed in panel B. Threshold line is vertical at *dτ* = 0 similar to figure 1 B.

Receptors are produced with fixed rate (rescaled to 1). Receptor-ligand complexes (*C_i_*) trigger response but are degraded with time-scale *τ_i_*, defining quality of ligands (see Figure 2 A). Steady state equations for homeostatic model is for mixture {(*L_n_,τ_n_*)}

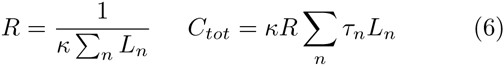

When only one type of ligand (*L*, *τ_c_*) is present, *C_tot_ = τ_c_* is adaptive (i.e. independent from *L*) as expected. At steady state, *R* is inversely proportional to total ligand concentration presented irrespective of their *τs*. Importantly, this is not due to a titration effect of a fixed pool: *R dynamically* buffers any ligand addition, so that steady state output *C_tot_* is a weighted linear combination of *τ_n_*s in 6. Such combination is always inferior to the maximum of *τ_n_*, so if such maximum indeed is *τ_c_*, antagonism ensues. If we consider the mixture of *L*_1_ ligands (critical time *τ_c_*) with *L_2_* ligands (*τ_c_* - *dτ*) we get

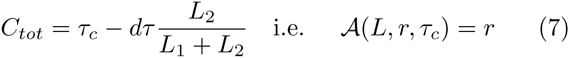

with *r = L_2_/L_2_+L_1_* as before. This contribution is exact and linear in *dτ* at all orders.

Figure 2B and C show values of *C_to_t* for pure ligands and mixture, with vertical discrimination lines at *τ_c_*. This simple model thus represents a perfect implementation of the principles sketched in Figure 1 [22]. Interestingly, antagonism strength is proportional to the deviation of antagonists’ quality *dτ* from *τ_c_* (equation 7). In particular, antagonism is very weak for ligand quality close to *τ_c_* and gets stronger as quality of ligands gets further below threshold.

This hierarchy of antagonism is completely opposite to what we know of immune examples, where antagonism is maximum for ligands close to *τ_c_*, and vanishes for small *τ_s_*, corresponding to “self [15]. Many immune models are based on kinetic proofreading [10, 18], proposed to be mediated by kinetic segregation [23]. Receptors exist in different phosphorylation states (*C_j_s*, Figure 3 A). Transition rates between states are functions of internal variables (notation M), accounting for all signalling inside cells (kinases, phosphatases), and assumed to diffuse freely and rapidly. M values depend on the total occupancy of some *C_j_s*. Downstream decision is usually assumed to be effectively controlled by thresholding on one state *C_n_R__*.

**FIG. 3.**
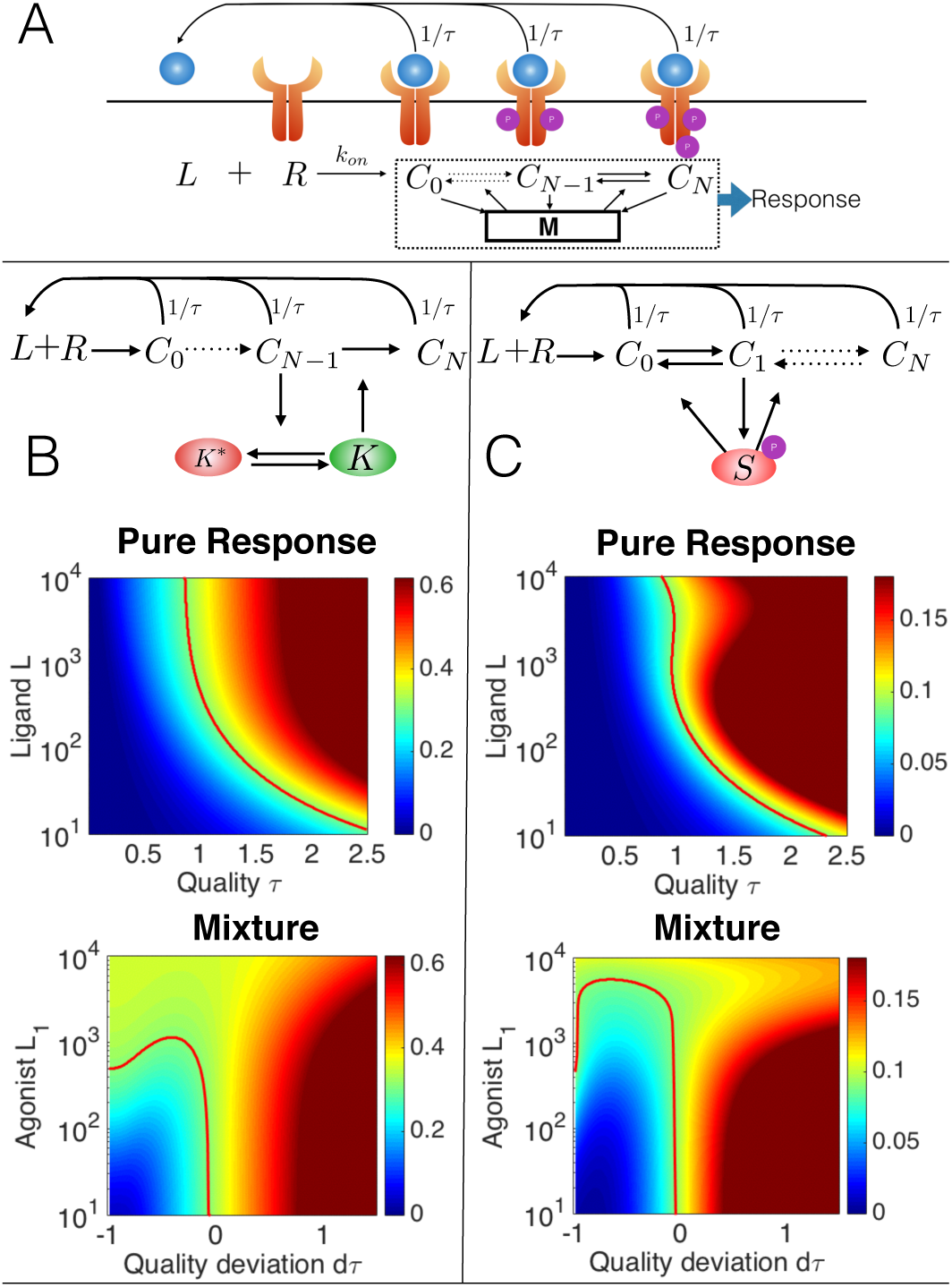
(A) General topology considered for proofreading-based models, with shared internal variables **M**. (B) Adaptive sorting topology and response, with colormap for output *C_n_* for pure ligands and mixtures. Equations and parameters are in Appendix. We fix *L*_2_ = 10^3^ ligands for mixtures, using similar conventions to Figure 2C. Threshold Θ for discrimination line was chosen so that response is triggered by *L*_1_ ∼ 500 ligands at *τ_c_*. For *dτ* ∼ − 1, downward folding of both response line and green region compared to Figure 2 C indicates decreased antagonism. (C) Immune model with same conventions as (B), equations and parameters are in Appendix

Using standard assumptions, it is easy to show (Appendix) that for this class of models, the number of receptor states *Cj* bound to ligands (*L*, *τ*) is:

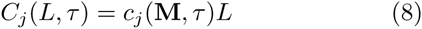

i.e. we can decouple influence of ligands into a linear contribution (*L*), and an indirect one (*c_j_*) purely due to internal variables. In the context of absolute discrimination equation 1 imposes that *C_n_R__* is tuned to threshold Θ irrespective of *L* at *τ_c_*. Equation 8 thus constrains *C_n_R__* to be inversely proportional to *L* at *τ_c_*, very much like *R* in homeostatic model. Thus internal variables necessary implement an incoherent feedforward loop [24] and/or a negative feedback via *c_n__R_*. This is observed in adaptive sorting models for high ligands *L* [5] (vertical asymptotic discrimination line on Figure 3 B). Immune model from [20] only locally approximates this adaptive constraint (Figure 3 C, high values of *L*).

An important difference with homeostatic model is that transition rates between variables depend explicitly on *τ* in proofreading models. It is informative to consider an Ansatz with only one internal variable *M* and separable influence of *τ* so that the contribution to the output variable from ligands (*L*, *τ*) is

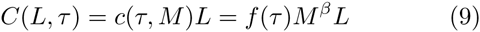

with 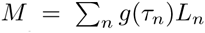 at steady state for mixture {*L_1_,τ_i_*}. *β* = −1 gives perfect adaptation but different values of *β* can give realistic biological effects such as loss of response at high ligand concentration as observed in [20]. Function *f*(*τ*) typically corresponds to the “direct” influence due to the main proofreading cascade (*n_R_* steps), while *g* encodes internal effects triggered by another proofreading step (*m*). Linear contribution to antagonism can be directly computed. We Taylor expand *C_tot_*({(1 − *r*)*L*,*τ_c_*;*rL*,*τ_c_* + *dτ*})−*C*(*L*,*τ_c_*) to get the linear term at order d*τ*:

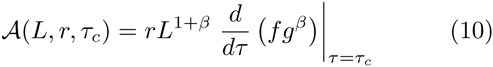

For β *=-1*, this term is very similar to antagonism expression for homeostatic model (equation 7), with extra *τ* dependency coming from f, *g*. If we impose that mixture of agonists trigger response (equation 2), we necessary have 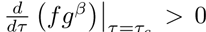. If 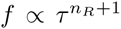 and *g* ∝ *τ*^m+1^ which is approximately the case for adaptive sorting (see Appendix), we thus have n_R_ + 1+β(m + 1) > 0, and thus for β *=* −1 we get *n_R_ > m*. In many immune models [5, 15, 20] this constraint is naturally realized because the internal variable is regulated by a much earlier step (m) than the output (*n_R_*) within the same proofreading cascade.

To understand what happens for *τ* << *τ_c_*, it is more illuminating to compute ratio of responses

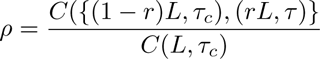

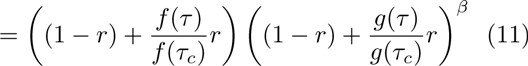

In models of immune detection based on proofreading such as adaptive sorting, *f* and *g* are powers of *τ*, so that *f*(*τ*)/*f*(*τ_c_*),*g*(*τ*)/*g*(*τ_c_*) << 1 when *τ* < *τ_c_*. So if β *=* −1, we see that *ρ →* + 1 when *τ*/*τ_c_ -+* 0, explaining why antagonism disappears in this parameter regime (corresponding to self in immune detection). Note that for *f* ∞ *τ*^*nR*+1^ and *g* ∝ *τ*^*m*+1^, the leading order correction at small *τ* is coming from the *g* term if *n_R_* > *m*, illustrated on Figure 4. Proofreading models interpolate between no antagonism for *τ* → 0 and linear antagonism close to threshold. Thus antagonism strength increases as *τ* increases from 0, as observed experimentally [15], before quickly collapsing again very close to threshold. Quality of ligands *τ* for maximum antagonism is closer to threshold *τ_c_* with increasing *m* and *n_R_ - m* (Figure 4).

**FIG. 4.**
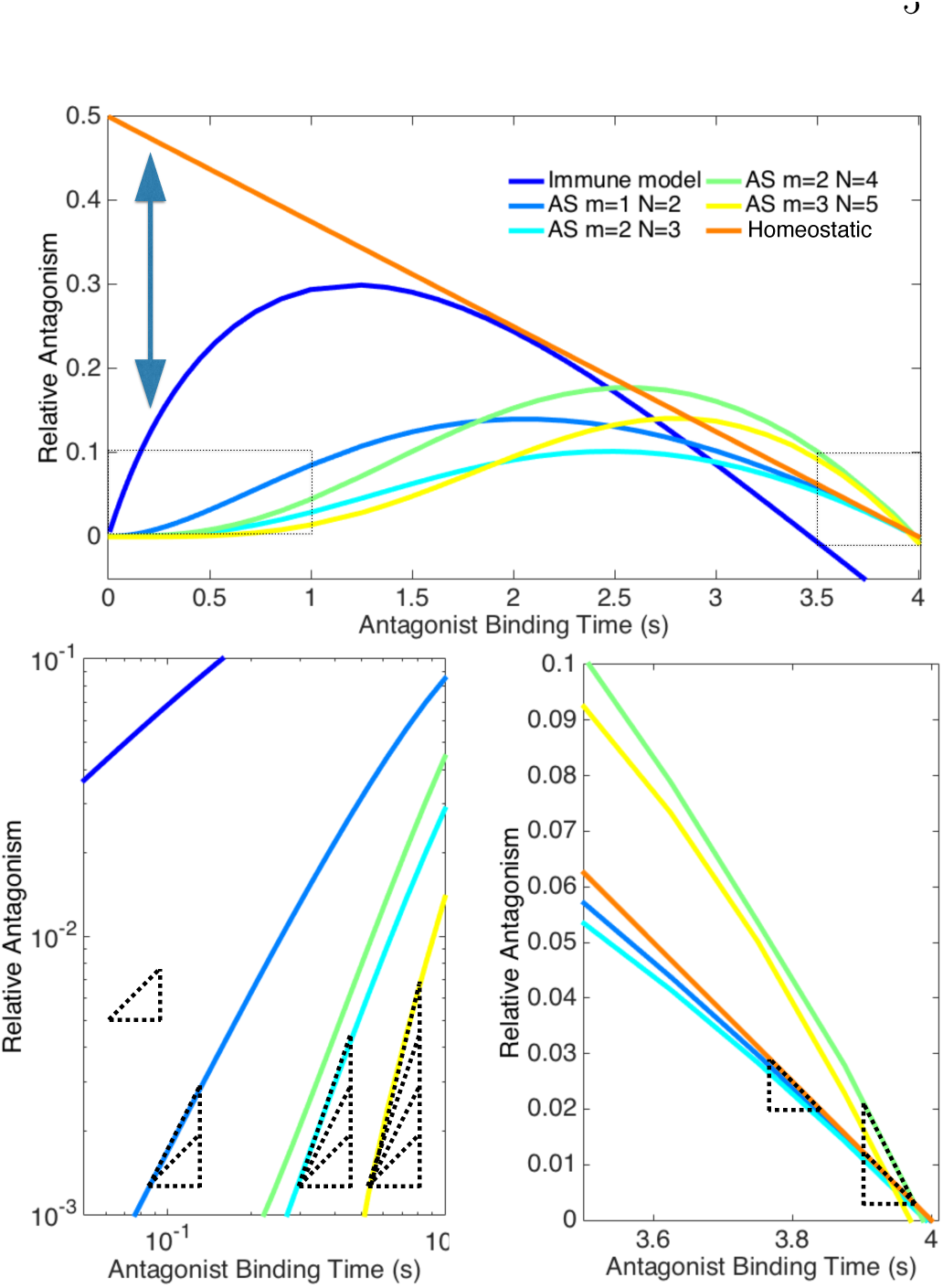
(Top) Computation of relative antagonism strength for different models with varying parameters. We consider *L_1_ = L_2_ =* 4 × 10 ligands, *τ_c_* = 4, and plot 1 − *C*{(*L*_1_,*τ_c_*), (*L*_2_,*τ*)}/*C*(*L*_1_,*τ_c_*). For instance, for homeo-static model, antagonism is 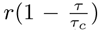. Double arrow indicates region of inverted hierarchy for antagonism. Dotted squared highlight regions plotted on bottom panels. AS indicates adaptive sorting model similar to Figure 3 B, Immune model corresponds to model of Figure 3 C. (Bottom left). Log-log plot, showing the dependency of antagonism for small *τ*. Dashed triangles indicate slopes of 1,2,3,4. For adaptive sorting models, slope is roughly equal to *m* + 1 as expected from the form of *g*. (Bottom right) Linear plot close to *τ = τ_c_*. Aspect ratio has been chosen so that slope for the homeostatic model *−r*/*τ_c_* is plotted with slope −1. Dashed triangles indicate slopes of −1,−2. For adaptive sorting models, slopes are thus roughly equal to −(*N* − *m*)*r*/*τ_c_* as expected from the Ansatz.

In conclusion, we have established the fundamental connection between ligand antagonism and absolute discrimination, the latter being a necessary by-product of the former, thus qualifying as a “phenotypic spandrel”. While there have been recent studies of spandrels at the level of protein structure [25], to our knowledge, this is the first study of an explicit spandrel structuring a complex signalling phenotype. With the exception of [26], our modelling hypotheses correspond to most phenotypic models of ligand discrimination we are aware of [10], as well as more complex models such as the one in [15]. Noise can be included as in [5, 20] via time-integration of some output variable, replacing all concentrations by expectation values, e.g. *C* → < *C* >. To possibly disentangle antagonism from absolute discrimination one would need to complexify formalism or hypotheses: for instance the time-course of a variable far from steady-state [26], coupled to multi stability [27] might be used. Discrimination might not be fully absolute and there might be additional modulations by ligand concentration or association rates [10, 28]. However if *τ* is the main discriminant for ligand quality as often hypothesized, this should only yield corrections to the leading order behaviour studied here.

The principles described can be applied to many pathways. We have already mentioned early recognition by T cells [15, 20]. Another relevant immune example are Fc receptors, displaying similar hierarchy of binding time sensitivity, and an explicit mechanism for antagonism [14] where antagonists titrate the kinase responsible for proofreading steps, therefore realizing exactly the adaptive sorting network topology [5]. Antagonism mediated by similar feed-forward/feedback mechanisms has been described in other contexts, such as Hh signalling [29]. Hormonal pathways present response curves very reminiscent of a model for immune recognition [20], such as non-monotonic response activity with varying ligand concentration [19], that would correspond in our framework to approximate adaptation. It has been established that in such a context antagonists differ from agonists purely based on slower binding kinetics [30], further suggesting an absolute discrimination mechanism. A recent meta-study shows that most of these pathways indeed use internal negative feedbacks to change monotonicity of response [31] (possibly leading to endocrine disruption for non natural ligands). Other qualitative recognition processes may present similar properties: for instance antagonism is observed in olfaction [32]. Clearly more studies are needed to quantify both absolute quality of ligands and antagonism, but all these examples are possibly structured by similar spandrels as described here.

Is antagonism an evolutionary spandrel related to immune detection ? Networks evolved in [5] systematically show local biochemical adaptation and antagonism, as expected from the present derivation. So evolutionary spandrels can be observed and studied in evolutionary simulations, and their emergence studied theoretically. In nature, it is difficult to definitely know if a trait is a spandrel without detailed historical data and comparisons [33]. Other possibilities could be that absolute discrimination has been selected in conjunction with other properties (such as information transmission [34, 35] or statistical decision [36]). But fundamentally, the presence of phenotypic spandrel here is due to the non-trivial computation performed by the cell to disentangle two parameters naturally tied by standard ligand-receptor interactions: kinetic of binding and ligand concentration. Experiments probing internal feedbacks (such as [14]) connect spandrels to cellular computations, similar to symmetries connected to physical laws.

## Acknowledgments

We thank Audrey H Moores, Jean-Benoît Lalanne and Grégoire Altan-Bonnet for comments and discussions. This project has been funded by the Natural Science and Engineering Research Council of Canada (NSERC), Discovery Grant Program, and a Simons Investigator Award for Mathematical Modelling of Biological Systems to PF. LS was supported by an Undergraduate Summer Research Award from NSERC.

## APPENDIX

### A. Equations for adaptive sorting (AS)

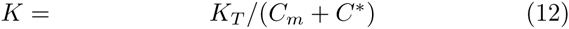

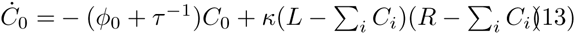

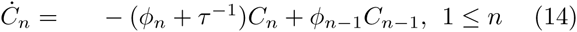

where *φ_m_ = φ_K_K*,*φ_N_ = 0* and *φ_n_ = φ* for other values of *n*.

Parameters used for simulation in main text are *κ =* 10^−5^, *R* = 3 x 10^4^,*φ* = *φK_T_ =* 0.09, *C*^*^ = 3, *m* = 2, *N* = 3. Output is *C_N_*. We used *τ_c_* = 4 *s* and defined threshold Θ = 0.31 to plot a discrimination line.

We get immediately at steady state

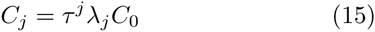

setting 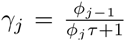 and 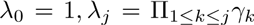. Neglecting 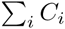 in front of *R* we have

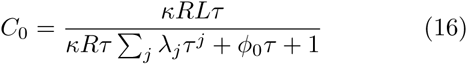

Assuming *φ_j_τ,κRτ* << 1, we then get the following scaling laws for *n ≤ m*

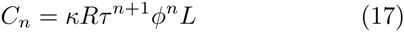

and for *n > m*

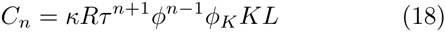

Since 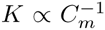, we recover scaling law of the Ansatz from the main text.

### B. Equations for immune model

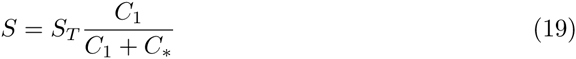

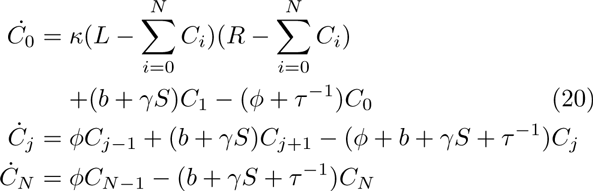

Parameters used for simulation in main text are *κ =* 10^−4^, *R =* 3 × 10^4^, *φ =* 0.09, *b* = 0.04, γ*S_T_* = 0.72, *C∗ =* 300. Output is *C_N_*. We used *τ_c_* = 4 *s* and defined threshold Θ = 0.09 to plot a discrimination line.

### C. Origin of linearity

Considering models defined above, it is clear that if we assume unsaturated receptors, *i.e*. 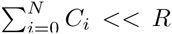, calling **C** the vector of occupancies and **M** the internal variables, we have:

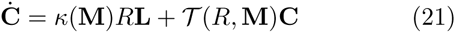

**L** = (*L*, 0,…, 0) is the vector corresponding to ligand input, *κ*(**M**) the association rate of ligands to receptors - by definition here ligands and receptors bind into the first state *C*_0_. *T*(*R*,**M**) is a matrix defining linear rates between occupancies states, depending on internal variables and parameter *τ*. Dynamics and steady state value of **M** is given by extra equations that are model specific (e.g. equations 12 and 19 above).

For such systems, we have at steady state

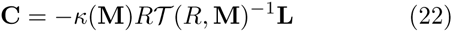

from which we can directly compute the *c_j_* as a function of **M**. Then we can use equations defining steady-state values of **M** as function of **C** to close the system.

When several ligand types are present, independence of ligand binding means that a system similar to equation 21 holds for every single vector of occupancy 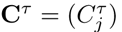 of receptor states bound to ligands with quality *τ*. Coupling between different types of ligands only occur via internal variables **M**. Note that we can also generalize this formalism so that transition rates depend on occupancies (i.e. effectively giving non linear transition rates between states) by assuming that internal variables **M** are occupancies themselves. The underlying strong assumption here is that the coupling is global, via total occupancies.

